# Investigation of the genus *Flavobacterium* as a reservoir for fish-pathogenic bacterial species: the case of *Flavobacterium collinsii*

**DOI:** 10.1101/2022.09.27.509832

**Authors:** Bo-Hyung Lee, Pierre Nicolas, Izzet Burcin Saticioglu, Benjamin Fradet, Jean-François Bernardet, Dimitri Rigaudeau, Tatiana Rochat, Eric Duchaud

## Abstract

Bacteria of the genus *Flavobacterium* are recovered from a large variety of environments. Among the described species, *Flavobacterium psychrophilum* and *Flavobacterium columnare* are causing considerable losses in fish farms. Alongside these well-known fish-pathogenic species, isolates belonging to the same genus recovered from diseased or apparently healthy wild, feral, and farmed fish have been suspected to be pathogenic. Here, we report the identification and genomic characterization of a *F. collinsii* isolate (TRV642) retrieved from rainbow trout spleen. A phylogenetic tree of the genus built by aligning the core genome of 195 *Flavobacterium* species revealed that *F. collinsii* is standing within a cluster of species associated to diseased fish, the closest one being *F. tructae* which was recently confirmed as pathogenic. We evaluated the pathogenicity of *F. collinsii* TRV642 as well as of *F. bernardetii* F-372^T^, another recently described species reported as a possible emerging pathogen. Following intramuscular injection challenges in rainbow trout, no clinical signs nor mortalities were observed. However, *F. collinsii* was isolated from the internal organs of wounded fish, suggesting that the bacterium could invade fish under compromised conditions such as stress and/or wounds. Our results suggest that some fish-associated *Flavobacterium* species should be considered as opportunistic fish pathogens causing disease under specific circumstances.

**IMPORTANCE:** Aquaculture has expanded significantly worldwide in the last decades and accounts for half of human fish consumption. However, infectious fish diseases are a major bottleneck for its sustainable development and an increasing number of bacterial species from diseased fish raise a great concern. The current study revealed phylogenetic associations with ecological niches among the *Flavobacterium* species. We also focused on *Flavobacterium collinsii* that belongs to a group of putative pathogenic species. The genome contents revealed a versatile metabolic repertoire suggesting the use of diverse nutrient sources, a characteristic of saprophytic or commensal bacteria. In a rainbow trout experimental challenge, the bacterium colonized only oppressed fish facing stressful conditions suggesting opportunistic pathogenic behavior. This study highlights the importance of experimentally evaluating the pathogenicity of the numerous bacterial species retrieved from diseased fish.

## INTRODUCTION

Members of the genus *Flavobacterium* (phylum *Bacteroidota*, family *Flavobacteriaceae*) are most frequently isolated from environmental sources. They are common in freshwater environments and in soil, especially in the rhizosphere, but strains have also been isolated from brackish or sea water and glaciers in temperate, tropical or polar areas (1). Following new molecular omics-based approaches, the taxonomy has been clarified and the number of formally described *Flavobacterium* species has rapidly expanded to include 268 species at the time of writing (lpsn.dsmz.de/genus/Flavobacterium accessed august 2022). The vast majority of these species are considered harmless, consuming inert organic matter thus playing an important role in biogeochemical cycles (2). Among them, *F. johnsoniae* has emerged as a model organism for studying gliding motility and protein secretion (3). In addition, the genus encompasses two very important fish pathogens, *F. psychrophilum* and *F. columnare*, both reported to cause considerable losses in farmed and wild freshwater fish. To account for its genomic and phenotypic diversity, the latter has been recently divided into four distinct species which have yet to be validated, namely, *F. columnare*, “*F. covae*”, “*F. davisii*” and “*F. oreochromis*” based on genomic comparisons as well as differences in host fish species and virulence (4). Another species, *F. branchiophilum*, is also known as a fish pathogen, but within more restricted geographical areas. In addition, the following species, often represented by a very small number of isolates, were recovered from diseased fish tissues and have been suspected to be pathogenic: *F. araucananum* from kidney and external lesions of Atlantic salmon (*Salmo salar*) (5); *F. bernardetii* from kidney and liver of rainbow trout (*Oncorhynchus mykiss*) (6); *F. turcicum* and *F. kayseriense* from rainbow trout kidney and spleen, respectively (7); *F. branchiarum* and *F. branchiicola* from rainbow trout gills (8); *F. chilense* from external lesions of rainbow trout (5); *F. collinsii* from the liver of rainbow trout (8); *F. hydatis* from the gills of diseased salmon (9, 10); *F. inkyongense* from diseased chocolate cichlids (*Hypselecara coryphaenoides)* (11); *F. johnsoniae*-like isolates from various diseased fish species (12); *F. oncorhynchi* from liver and gills of rainbow trout (13); *F. piscis* from liver, gills, and kidney of rainbow trout (14); *F. plurextorum* from liver and eggs of rainbow trout (15); and *F. succinicans* from rainbow trout gills suffering bacterial gill disease (16). At least one species, *F. tructae*, which was isolated from liver, gills, and kidney of rainbow trout (14) and concurrently from kidney of feral spawning adult Chinook salmon (*Oncorhynchus tshawytscha*) under the alternative name of *F. spartansii* (17), may be considered a salmonid pathogen as two isolates were able to induce pathological changes and mortality in experimentally infected Chinook salmon, though only using very high infectious doses (18). In contrast, most of the aforementioned fish-associated species have not been assessed for their level of virulence using experimental challenges.

The lack of information about the pathogenicity on many *Flavobacterium* species can lead to unnecessary and irrational use of antimicrobials, and, on the opposite, to the lack of surveillance of bacteria that can cause disease outbreaks. Interestingly, the continuous increase in whole genome sequences generate data that may help identifying ecological niches and pathogenicity, by providing improved phylogenetic resolution and access to gene repertoires underpinning the phenotypes. As an initiative for this approach in the genus *Flavobacterium*, we exploited the large number of genomes available to investigate the relationship between phylogeny and environmental niches at the species level. In parallel and in attempt to satisfy Koch’s postulates, we assessed the virulence of two recent isolates of *F. bernardetii* (6) and *F. collinsii*, a species closely related to *F. tructae*, using experimental infection in rainbow trout.

## MATERIALS AND METHODS

### Bacterial strains and growth conditions

Bacteria were grown on tryptone yeast extract salts (TYES) broth [0.4% w/v tryptone, 0.04% w/v yeast extract, 0.05% w/v MgSO_4_ 7H_2_O, 0.02% w/v CaCl_2_ 2H_2_O, 0.05% w/v D-glucose, pH 7.2] or on TYES agar (TYESA) supplemented with 5% fetal bovine serum for 4 days at 18°C. Precultures were prepared using a single colony in TYES broth and grown overnight at 18°C and 200 rpm. Cultures were prepared by dilution of stationary preculture and grown overnight to the early stationary phase (OD600 of 1.0 ± 0.1) for infection challenges. Cultures were 10-fold serially diluted in 1% w/v peptone water and colony-forming units (CFUs) were counted *a posteriori* after 48-72 h of incubation at 18°C on TYESA. For long term storage stationary bacterial cultures were stored at −80°C with 20% v/v glycerol.

### 16S rRNA gene sequencing

PCR amplification was performed using a universal bacterial 16S rRNA primer set composed of forward (27F: 5’-AGA GTT TGA TCM TGG CTC AG-3’) and reverse (1492R: 5’-TAC GGY TAC CTT GTT ACG ACT T-3’) primers. The PCR mixture contained final concentrations of 0.3 μM primers each, 1.25 Units of Dream Taq DNA Polymerase and 1X buffer (Thermo Fisher Scientific), 0.3 mM dNTPs and 1.5 μl bacterial DNA prepared by heat lysis (at 99°C for 10 min) of stationary culture. The volume was adjusted to 50 μl with ddH20 and PCR was performed with initial denaturation at 95°C for 5 min followed by 30 cycles of amplification [95°C for 20 s for denaturation, 48°C for 20 s for annealing, 72°C for 2 min for extension], and finished by extension at 72°C for 10 min. PCR amplicons were sequenced by Sanger sequencing.

### Rainbow trout challenges

The rainbow trout synthetic line (*O. mykiss* Sy) selected by INRAE (19) was used for experimental infections and median lethal dose (LD50) estimation. Experiments were carried out independently for *F. bernardetii* F-372^T^, *F. collinsii* TRV642 and *F. psychrophilum* FRGDSA 1882/11 using rainbow trout fingerlings weighing 6.3g, 6.7g and 8.4g on average, respectively. Fish were reared at 10°C in recirculating aquaculture system (RAS) in 30-liter tanks with dechlorinated tap water. Two days before infection, 7 groups of 10 fish for each bacterial strain were transferred to the biosafety level 2 (BSL2) zone in 15-liter tanks with flow water (1 renewal per hour). Six groups were challenged at different doses, as follows: 6 suspensions were prepared by 10-fold serial dilutions (10^−1^ to 10^−6^) of a bacterial culture, then 50 μl of each suspension were administered per fish by intramuscular injection at a point midway between the insertion of the dorsal fin and the lateral line following anesthesia (50 mg/L, MS222, Sigma-Aldrich). For *F. psychrophilum*, only 5 groups corresponding to 10^−3^ to 10^−7^ dilutions were used and the median lethal dose was estimated using the moving average and interpolation method described by Thompson (20) with 3 dose-groups used to calculate each moving average. In each challenge, one group of 10 fish was injected with sterile TYES broth as a negative control. All fish were maintained using flow water at 10°C. Mortality was recorded twice a day and fish were monitored for clinical signs and behavioral changes throughout the challenge. Euthanasia was carried out by bathing fish in tricaine at concentration of 300 mg/L (MS222, Sigma-Aldrich). Dead or euthanized fish were examined for the presence of bacteria in organs including the spleen, kidney and liver. Organs were homogenized in peptone water containing silica spheres using a FastPrep instrument (MP Biomedicals) at 6 m/s for 20 sec. Tissue homogenates were streaked on TYESA and kept for 3-5 days at 18°C before examination for bacterial growth.

### Sequencing and annotation of the *F. collinsii* genome

Genomic DNA (gDNA) was extracted from stationary broth culture using genomic DNA-Tip 100/G system and buffer set (Qiagen) following the manufacturer’s instruction. For short read sequencing, the library was constructed using TruSeq genomic kit (Illumina) and paired-end sequenced on a NextSeq instrument using NextSeq 500/550 Mid Output Kit v2 (Illumina). In parallel, gDNA was sequenced on GridION (Oxford Nanopore) using a FLO-MIN106 flowcell for long-read sequencing. Hybrid genome assembly was performed using Unicycler v0.4.8 available on the PATRIC platform with default parameters (21). Genome annotation and comparison were performed using the MicroScope platform (22). InterProScan was used to identify susCD pairs using IPR023996 and IPR023997 for SusC and IPR012944, IPR024302, IPR041662, IPR033985 for SusD with co-localization criteria. Proteins secreted by the T9SS harboring a conserved C-terminal domains (CTDs) were identified using IPR026444 and IPR026341 for type A and B CTDs (23), respectively. CAZymes were identified using dbCAN, which combines three tools; hits found by only one tool were removed to improve accuracy (24). Peptidases were identified using the MEROPS database (https://www.ebi.ac.uk/merops/) (25).

### Phylogenetic analysis

Average nucleotide identities (ANI) were computed using OrthoANIu and the web-service http://www.ezbiocloud.net/tools/ani (26).

Genome records were retrieved from the RefSeq database (https://www.ncbi.nlm.nih.gov/datasets/genomes/?taxon=237&utm_source=data-hub, accessed in April 2022) (27).

Proteomes were compared pairwise with Blastp (v2.12, low-complexity filter disactivated, e-value cut-off 1e-5, otherwise default settings) (28) and the results served for a single linkage clustering of the genes based on a criterion of 45% amino-acid sequence identity over 70% of the gene length. Conserved single copy genes were identified as clusters with a single representative in each genome and their sequences subjected to multiple sequence alignment using Muscle (v3.8.31, default setting) (29).

Neighbor-Joining phylogenetic tree reconstruction was conducted using FastTree (v2.1.10, default settings) (30) on the concatenated multiple sequence alignments, after alignment gaps removal. Statistical support of the tree topology was assessed by comparison with trees reconstructed on 400 bootstrap replicates of the original alignment.

## RESULTS

### A new *F. collinsii* isolate retrieved from rainbow trout

In order to survey the health status of rainbow trout raised at INRAE fish facilities (IERP), different fish organs of dead animals are regularly sampled to perform basic bacteriological quality controls. Organs (*i.e*., usually spleen and liver) are crushed in TYES broth and the lysates are spread on Petri dishes to ensure the absence of any pathogenic *Flavobacterium* species. Surprisingly, during the summer 2020, yellow-pigmented bacterial colonies with high spreading activity were observed after streaking the spleen of one rainbow trout fingerling (2 g) that had died without any disease symptom. The fish belonged to the highly *F. psychrophilum*-susceptible A36 isogenic line (31). A pure culture was obtained from one bacterial colony and 16S rDNA sequencing identified the bacterium as probably belonging to the species *F. collinsii*. For a more accurate characterization, the complete genome of this isolate, named TRV642, was resolved by high-throughput sequencing using the hybrid assembly of long Nanopore reads and short Illumina reads. The ANI between strain TRV642 and the *F. collinsii* type strain CECT 7796^T^ was 98.30%, far above the threshold delineation (cutoff 95–96%) of a bacterial species (32), thus confirming the initial 16S rDNA taxonomic affiliation.

### Phylogenetic analysis and correlation with the ecological niche at the genus level

To cover the diversity of the species with complete genome records available in public databases, we extracted from the RefSeq database all genomes belonging to genus *Flavobacterium* with the tag “representative”, constituting a set of 195 genomes from distinct species. Gene repertoire comparisons delineated a core of 510 conserved single copy genes. Their multiple sequence alignment after gap removal represented 154,805 amino-acids (98,363 polymorphic sites) and was used for phylogenetic reconstruction. A tentative phylogenetic tree for the 195 *Flavobacterium* species is shown in Figure 1. The genome size and G+C-content to the corresponding medians for the genus were also compared (Figure 1).

**Fig 1.**
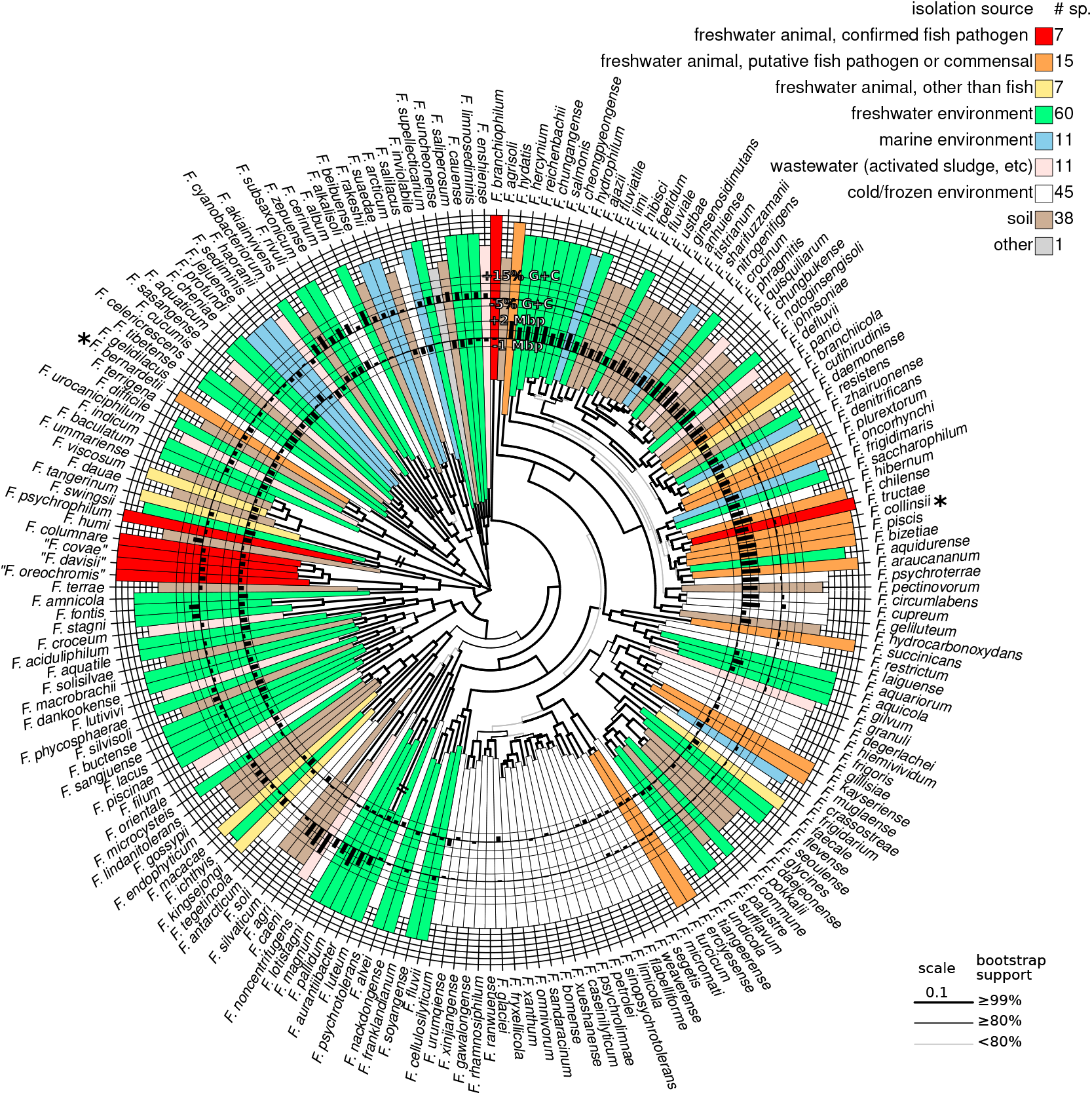
Tentative phylogenetic tree of 195 *Flavobacterium* species. Tree based on the protein sequences of the 510 single-copy genes of all *Flavobacterium* species which complete genome is available in public databases. For clarity, the tree was midpoint rooted. Genome size and G+C contents are compared to the medians for genus and shown as circular barplots. Color and position on outer circle indicate the source of isolation of the type strain with confirmed pathogen species in red and outermost position.

To examine the links between position in the phylogenetic tree and ecological niche we retrieved from the original publication describing each species the environment from which the type strain was sampled. The diversity of environments was summarized into 9 different categories (Supplemental data set 1 and Figure 1), distinguishing 3 categories for the 29 species isolated from animal-associated samples, 5 categories for the 165 species isolated from other important natural environments, and 1 category “other” to account for the single species isolated in a totally different and more artificial context (*F. supellecticarium* isolated from a “synthetic wooden board”).

According to this classification, the 165 species (84.6% of the total) originating from non-animal related environmental samples were composed of 60 species (30.8%) retrieved from freshwater, 45 species (23.1%) from cold/frozen environments (predominantly Antarctic soil or glacier), 38 species (19.5%) from soils, 11 species (5.6%) species from marine environments (including sea sediments and algae), and 11 species (5.6%) from wastewater-related environments. Out of the 29 species (14.9% of the total) retrieved from, or in contact with, animals, only 7 of these animals were not fish. The 22 fish-associated species (11.3% of the total) were further decomposed into 7 confirmed pathogens (4 of which were previously considered as genomovars of *F. columnare*) and 15 putative pathogens or fish commensals. Taken together, the classification of the environments and the phylogenetic tree shed light on the possible connections between evolutionary links and ecological niches in the genus *Flavobacterium*. Among the patterns that emerge from this picture, a group around *F. glaciei* is mostly composed of species retrieved from cold/frozen environments. Some links between phylogenetic position and genome characteristics were also observed. In terms of G+C content, 8 species with particularly high G+C form a phylogenetic group around *F. caeni*. In terms of genome size, a large phylogenetic group encompassing more than a quarter of all considered species (51 species, 26.2%) including *F. johnsoniae* displays genomes up to 2 Mbp larger than the median value for the genus.

Whatever the specific category considered (“fish pathogen”, “fish-associated”, or “animal other than fish”), species associated with animals are scattered across the phylogenetic tree. Exceptions are the 4 fish-pathogenic species resulting from the splitting of *F. columnare* which group together, 2 pairs of closely related “fish-related species” (resp. *F. plurextorum*-*F. oncorhynchi* and *F. erciyesense*-*F. turcicum*) and a more important cluster of species around *F. tructae* that includes *F. collinsii*.

### *F. collinsii* belongs to a cluster of fish-associated species with large genomes

Strikingly, *F. collinsii* is the closest relative of the confirmed pathogen *F. tructae* but the two species are well distinct as confirmed by the ANI of only 87.56% between *F. collinsii* TRV642 and *F. tructae* MSU. These two species belong to a cluster of putative pathogenic species with *F. araucananum, F. bizetiae, F. piscis, and F. chilense*. All these species were isolated from diseased salmonid fish except for *F. bizetiae* whose origin of isolation was more loosely described as “a diseased freshwater fish” (33). All these species are predicted to derive from a common ancestor, also shared with *F. aquidurense, F. psychroterrae* and *F. hibernum*, 3 species not retrieved from animal-related material. This cluster, enriched in fish-related species comprising *F. collinsii* and *F. tructae*, belongs to the above-mentioned large genome group comprising *F. johnsoniae*.

The complete genome of *F. collinsii* TRV642 consists of a circular chromosome of 5,554,530 bp and predicted to contain 4,285 CDS, 69 tRNA genes, and 7 rRNA operons. The genome size is slightly smaller (−181,002 bp) than that of the *F. collinsii* type strain CECT 7796^T^. All species belonging to the aforementioned cluster comprising *F. tructae* also possess large genomes (from 5.4 to 6.1 Mbp), comparable to the 6.1 Mbp genome of the well-studied environmental species *F. johnsoniae* (34). These genomes are about 2-fold larger than the 2.9 Mbp genome of *F. psychrophilum* (35) and the 3.2 Mbp genome of the *F. columnare* type strain ATCC 23463^T^ (4), the two well-characterized and unquestionably fish-pathogenic species of the genus.

### Insights from the gene repertoire: protein secretion

The type IX secretion system (T9SS) is responsible for protein secretion and required for gliding motility (3). This machinery is confined to the phylum *Bacteroidota* and genes encoding the core components of the T9SS machinery, the attachment complex, and the *gld* genes needed for gliding motility were all identified in the *F. collinsii* TRV642 genome (Supplemental data set 2).

Proteins secreted by the T9SS possess conserved C-terminal domains (CTDs). In the *F. collinsii* TRV642 genome, 45 genes encoding proteins with a type A (TIGR04183) CTD were identified (Supplemental data set 3), some of which with predicted enzymatic functions, as well as 9 genes encoding proteins with a type B (TIGR04131) CTD, none with predicted enzymatic function. Enzymes with a type A CTD include 9 peptidases likely involved in protein/peptide degradation, all conserved in the genomes of *F. tructae* and *F. chilense* but absent, with a few exceptions, in those of *F. psychrophilum* and *F. columnare*. Among the other type A CTD enzymes, two are polysaccharide lyases and one is a glycoside hydrolase likely involved in carbohydrate degradation. This repertoire of T9SS-secreted enzymes supports the view of *F. collinsii* as a degrader of high molecular weight organic matter and suggests metabolic versatility, a functional trait shared by some members of the family *Flavobacteriaceae*, such as *F. johnsoniae* (34).

In addition to the *Bacteroidota-*specific T9SS, a B-type T4SS is also present in the *F. collinsii* TRV642 genome (Supplemental data set 2). This versatile secretion system is utilized to mediate horizontal gene transfer and also allows Gram-negative pathogenic bacteria to translocate a wide variety of virulence factors into the host cell (36). Because the *F. collinsii* T4SS locus encompasses genes involved in DNA transfer and encoding relaxase/mobilization family proteins (*i.e*., MobA/VirD2 and MobC), it is tempting to speculate that *F. collinsii* T4SS is dedicated to the recruitment and delivery of DNA substrates.

### Insights from the gene repertoire: Polysaccharides Utilization Loci and CAZymes

Members of the phylum *Bacteroidota* have developed multi-component protein systems aimed at sensing, binding, transporting, and degrading specific glycans (37). Genes encoding these systems are often co-localized in regions referred to as Polysaccharide Utilization Loci (PULs). Tandem *susD*-like and *susC*-like genes, which encode a carbohydrate-binding lipoprotein and a TonB-dependent transporter, respectively, are considered a hallmark of PULs and these *susCD*-like gene pairs are generally used to identify PULs in *Bacteroidota* genomes (38). It has been suggested that *susCD*-like pairs could transport other substrates than carbohydrates (34) and this was recently confirmed by the finding that the BT2263–2264 pair of *B. thetaiotaomicron* as well as the RagAB transporter of *Porphyromonas gingivalis* import oligopeptides (39, 40). The *F. collinsii* TRV642 genome contains 29 *susCD*-like pairs, most of which are encompassed in *bona fide* PULs, namely with adjacent genes encoding obvious additional polysaccharide utilization proteins (*i.e*., carbohydrate-active enzymes or CAZymes) (Table 1; Supplemental figure). About half of these PULs have been previously identified in the *F. johnsoniae* UW101 genome (34). For instance, PUL TRV642_1394-1405 is predicted to be dedicated to α-glucans/starch degradation (corresponding to PUL Fjoh_1398-1408), PUL TRV642_1594-1602 to β-glucans/xylan degradation (corresponding to PUL Fjoh_1559-1567), and PUL TRV642_4202-4209 to chitin degradation (corresponding to PUL Fjoh_4555-4564). In addition to these PULs shared with *F. johnsoniae*, some are likely of functional importance. For instance, PUL TRV642_0198-0209 encompasses 5 peptidases (belonging to families S9, S37, M24 and M43) and is likely involved in the uptake of oligopeptides. PUL TRV642_4544-4556 encompasses 12 genes among which TRV642_4547 encoding a polysaccharide lyase (family PL8) protein precursor containing a type A T9SS C-terminal secretion signal likely anchored in the outer-membrane, facing the extracellular milieu, and *hepC* encoding a protein highly similar (63% aa identity) to heparinase III of the non-pathogenic soil bacterium *Pedobacter heparinus*, degrading heparan sulfate (41) and likely located in the periplasm. These two genes, organized in tandem, as well as other glycoside hydrolase (GH)-, polysaccharide lyase (PL)- and sulfatase-encoding genes are likely involved in the degradation of complex sulfated carbohydrate(s) belonging to the glycosaminoglycan(s) family such as heparan sulfate.

**Table 1.** *susCD* pairs identified in the *Flavobacterium collinsii* TRV642 genome.

| Locus_tags | Product | InterPro Id | In a *bona fide*<br>PUL | Predicted<br>substrate |
| --- | --- | --- | --- | --- |
| TRV642_0169 | SusC-like protein | IPR023996 and IPR023997 | No | Unknown |
| TRV642_0168 | SusD-like protein | IPR012944 and IPR033985 |  |  |
| TRV642_0202 | SusC-like protein | IPR023996 and IPR023997 | Yes | Oligopeptide |
| TRV642_0201 | SusD-like protein | IPR041662 |  |  |
| TRV642_0403 | SusC-like protein | IPR023996 and IPR023997 | No | Unknown |
| TRV642_0404 | SusD-like protein | IPR041662 |  |  |
| TRV642_0405 | SusC-like protein | IPR023996 and IPR023997 | No | Unknown |
| TRV642_0406 | SusD-like protein | IPR041662 |  |  |
| TRV642_0407 | SusC-like protein | IPR023996 and IPR023997 | No | Unknown |
| TRV642_0408 | SusD-like protein | IPR041662 |  |  |
| TRV642_0663 | SusC-like protein | IPR023996 and IPR023997 | No | Unknown |
| TRV642_0664 | SusD-like protein | IPR041662 |  |  |
| TRV642_0665 | SusC-like protein | IPR023996 and IPR023997 | No | Unknown |
| TRV642_0666 | SusD-like protein | IPR041662 |  |  |
| TRV642_1402 | SusC-like protein | IPR023996 and IPR023997 | Yes | α-glucans/starch |
| TRV642_1403 | SusD-like protein | IPR012944 and IPR033985 |  |  |
| TRV642_1595 | SusC-like protein | IPR023996 and IPR023997 | Yes | β-glucans/xylan |
| TRV642_1596 | SusD-like protein | IPR012944 and IPR033985 |  |  |
| TRV642_1932 | SusC-like protein | IPR023996 and IPR023997 | Yes | Oligosaccharides |
| TRV642_1933 | SusD-like protein | IPR041662 |  |  |
| TRV642_1968 | SusC-like protein | IPR023996 and IPR023997 | No | Unknown |
| TRV642_1969 | SusD-like protein | IPR012944 and IPR033985 |  |  |
| TRV642_2080 | SusC-like protein | IPR023996 and IPR023997 | No | Unknown |
| TRV642_2081 | SusD-like protein | IPR012944 and IPR033985 |  |  |
| TRV642_2255 | SusC-like protein | IPR023996 and IPR023997 | No | Unknown |
| TRV642_2254 | SusD-like protein | IPR033985 |  |  |
| TRV642_2325 | SusC-like protein | IPR023996 and IPR023997 | No | Unknown |
| TRV642_2326 | SusD-like protein | IPR041662 |  |  |
| TRV642_2567 | SusC-like protein | IPR023996 and IPR023997 | Yes | Oligosaccharides |
| TRV642_2566 | SusD-like protein | IPR012944 and IPR033985 |  |  |
| TRV642_2576 | SusC-like protein | IPR023996 and IPR023997 | No | Unknown |
| TRV642_2575 | SusD-like protein | IPR012944 and IPR033985 |  |  |
| TRV642_2869 | SusC-like protein | IPR023996 and IPR023997 | Yes | Oligosaccharides |
| TRV642_2870 | SusD-like protein | RagB/SusD domain |  |  |
| TRV642_3296 | SusC-like protein | IPR023996 and IPR023997 | No | Unknown |
| TRV642_3295 | SusD-like protein | IPR033985 |  |  |
| TRV642_3433 | SusC-like protein | IPR023996 and IPR023997 | No | Unknown |
| TRV642_3432 | SusD-like protein | IPR041662 |  |  |
| TRV642_3468 | SusC-like protein | IPR023996 and IPR023997 | Yes | α-glucans |
| TRV642_3467 | SusD-like protein | IPR012944 and IPR033985 |  |  |
| TRV642_3614 | SusC-like protein | IPR023996 and IPR023997 | No | Unknown |
| TRV642_3613 | SusD-like protein | IPR012944 and IPR033985 |  |  |
| TRV642_3711 | SusC-like protein | IPR023996 and IPR023997 | Yes | Oligosaccharides |
| TRV642_3712 | SusD-like protein | IPR012944 and IPR033985 |  |  |
| TRV642_3733 | SusC-like protein | IPR023996 and IPR023997 | Yes | Pectins |
| TRV642_3732 | SusD-like protein | IPR012944 and IPR033985 |  |  |
| TRV642_3861 | SusC-like protein | IPR023996 and IPR023997 | Yes | Oligopeptide |
| TRV642_3860 | SusD-like protein | IPR012944 and IPR033985 |  |  |
| TRV642_4207 | SusC-like protein | IPR023996 | Yes | Chitin |
| TRV642_4206 | SusD-like protein | IPR024302 |  |  |
| TRV642_4321 | SusC-like protein | IPR023996 and IPR023997 | No | Unknown |
| TRV642_4322 | SusD-like protein | IPR012944 and IPR033985 |  |  |
| TRV642_4333 | SusC-like protein | IPR023996 and IPR023997 | No | Unknown |
| TRV642_4334 | SusD-like protein | IPR012944 and IPR033985 |  |  |
| TRV642_4524 | SusC-like protein | IPR023996 and IPR023997 | Yes | Oligopeptide |
| TRV642_4525 | SusD-like protein | IPR012944 and IPR033985 |  |  |
| TRV642_4549 | SusC-like protein | IPR023996 and IPR023997 | Yes | Heparine/chondroitin sulfate |
| TRV642_4550 | SusD-like protein | IPR012944 and IPR033985 |  |  |

Genome analysis revealed a large gene repertoire (122) of carbohydrate-active enzyme (CAZymes) modules encompassed in 112 genes. Strikingly, this gene repertoire is approximately twice larger than in representative strains of pathogenic species *F. psychrophilum* (56) and *F. columnare* (63) (Table 2). Overrepresentation is even more obvious when only CAZymes dedicated to carbohydrate degradation are taken into account; for example, the *F. collinsii* genome encodes 51 GH and 9 PL. In contrast, the *F. psychrophilum* JIP02/86 genome contains 8 GH and 0 PL while that of *F. columnare* ATCC 49512 genome contains 13 PL and 3 PL. The high number of genes encoding CAZymes is a trait shared with the genomes of *F. tructae* and the cluster of putative pathogenic species (*i.e., F. collinsii, F. araucananum, F. bizetiae, F. piscis, and F. chilense*). These numbers range from 112 for *F. collinsii* to 238 for *F. piscis* (Table 2), the latter being comparable to the number found in the environmental bacterium *F. johnsoniae* (243). Therefore, data from T9SS secreted proteins, PUL systems and CAZymes indicate that *F. collinsii* evolved to take advantage of a large variety of both proteinaceous and carbohydrate substrates.

**Table 2.** Number of CAZymes modules encoded by *Flavobacterium* species.

| Species | Strain | Environment | Genome size (Mbp) | Glycoside hydrolases | Glycosyl transferases | Polysaccharide lyases modules | Carbohydrate esterases | Carbohydrate-binding | Total modules | Total genes |
| --- | --- | --- | --- | --- | --- | --- | --- | --- | --- | --- |
| *Flavobacterium johnsoniae* | UW101 | Fresh water, soil | 6.09 | 157 | 57 | 12 | 13 | 20 | 259 | 243 |
| *Flavobacterium psychrophilum* | JIP02/86 | Pathogenic essentially for *Salmonidae* | 2.86 | 8 | 44 | 0 | 4 | 2 | 58 | 56 |
| *Flavobacterium branchiophilum* | FL-15 | Pathogenic essentially for *Salmonidae* | 3.55 | 26 | 52 | 2 | 5 | 4 | 89 | 85 |
| *Flavobacterium columnare* | ATCC 49512 | Pathogenic for many freshwater fish species | 3.16 | 13 | 42 | 3 | 4 | 3 | 65 | 63 |
| *Flavobacterium collinsii* | TRV642 | Putative pathogen | 5.55 | 51 | 48 | 9 | 7 | 7 | 122 | 112 |
| *Flavobacterium chilense* | DSM 24724 | Putative pathogen | 6.11 | 124 | 45 | 14 | 11 | 22 | 216 | 198 |
| *Flavobacterium tructae* | MSU | Putative pathogen | 5.35 | 50 | 50 | 7 | 7 | 7 | 121 | 113 |
| *Flavobacterium piscis* | CCUG 60099 | Putative pathogen | 5.9 | 162 | 50 | 10 | 12 | 17 | 251 | 238 |
| *Flavobacterium bizetiae* | CIP 105534 | Putative pathogen | 5.87 | 149 | 50 | 23 | 13 | 17 | 252 | 222 |
| *Flavobacterium araucanum* | DSM 24704 | Putative pathogen | 6.01 | 108 | 48 | 7 | 8 | 20 | 191 | 174 |

### Insights from the gene repertoire: Antibiotic biosynthesis

The *F. collinsii* genome contains two large gene clusters (203 kb for TRV642_2161-2233 and 54 kb for TRV642_4252-4268) encoding non-ribosomal peptide/polyketide synthase enzymes likely involved in antibiotic biosynthesis. The genomes of *F. tructae* and *F. johnsoniae* also contain antibiotic biosynthesis gene clusters whereas those of *F. psychrophilum* and *F. columnare* are devoid of these loci, as well as of other antibiotic biosynthesis encoding genes.

### Virulence assessment in rainbow trout

With the increasing number of *Flavobacterium* species isolated from diseased fish worldwide (42), a better characterization of their lifestyle and their impact on fish health is needed. Within the time frame of this study, a novel *Flavobacterium* species named *F. bernardetii* was described after the recovery of two isolates from diseased rainbow trout exhibiting neurological symptoms (6). Together with *F. collinsii*, this species was proposed as a possible pathogen although pathogenicity was not properly validated using experimental infection.

In an attempt to evaluate the virulence of *F. collinsii* TRV642 and *F. bernardetii* F-372^T^ in rainbow trout, groups of fish were experimentally infected by intramuscular injection and maintained in flow water at 10°C for assessment of symptoms and mortality for two weeks. In this experimental infection model, the positive control strain FRGDSA 1881/11, belonging to the pathogenic species *F. psychrophilum*, was responsible for high mortality with a LD50 of 4.9 × 10^2^ CFU (Figure 2A).

**Fig 2.**
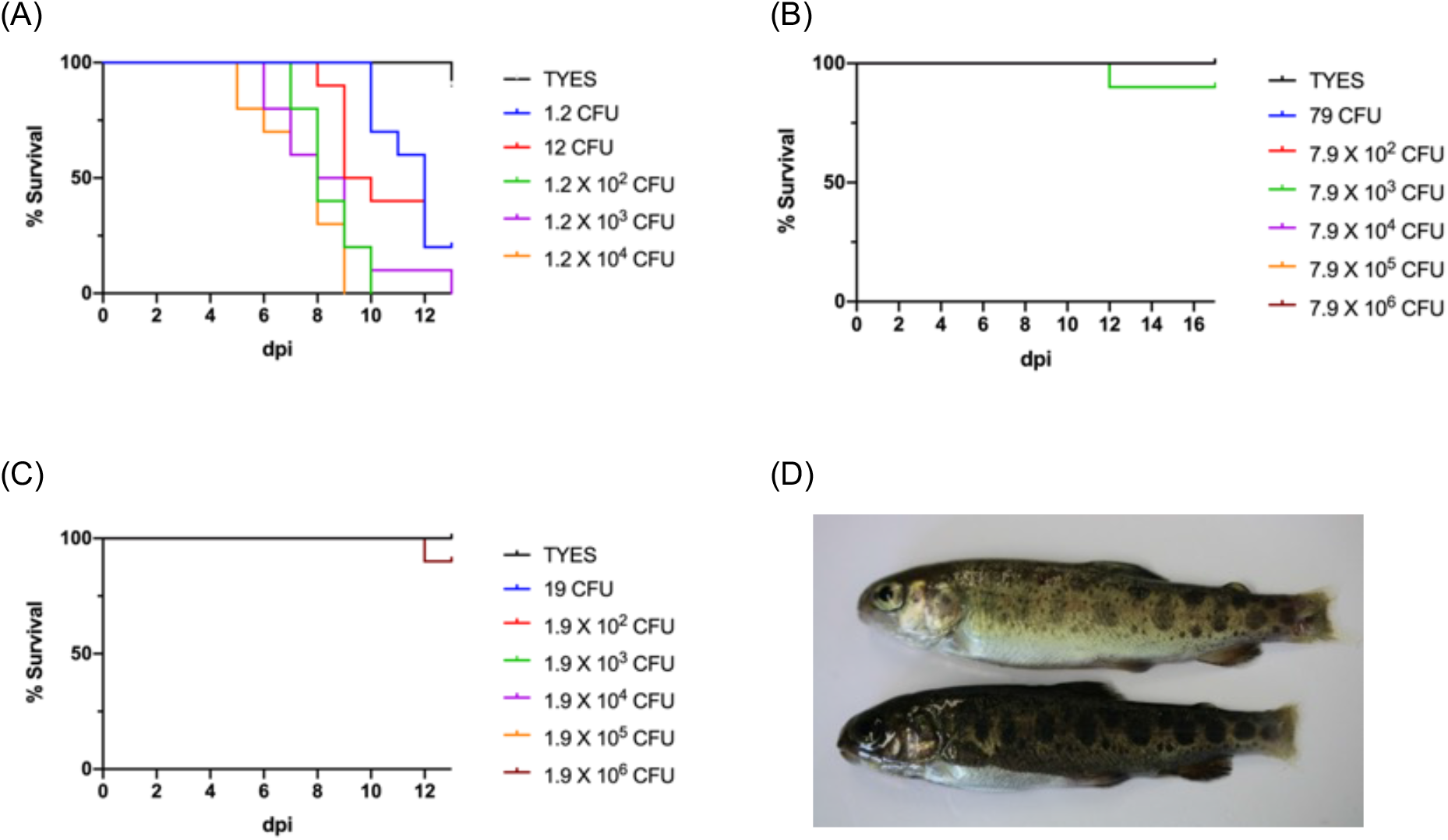
Evaluation of the virulence in rainbow trout. Kaplan-Meier survival curves of rainbow trout fingerling (Sy line, 6-8g) after intramuscular injection with (A) *F. psychrophilum* FRGDSA 1882/11, (B) *F. bernardetii* F-372^T^ and (C) *F. collinsii* TRV642 bacterial suspensions with doses ranging over six orders of magnitude. TYES broth was used as a negative control. (D) Fish infected with *F. collinsii* TRV642 and attacked by dominant fish display damaged caudal fin and tissue.

Only one fish injected with the medium infectious dose of *F. bernardetii* F-372^T^ (7.9 × 10^3^ CFU/fish) died at 12 dpi (Figure 2B) without any symptom and internal organs, including the brain, showed no trace of *F. bernardetii*. At 17 dpi, 3 fish belonging to the group that had received the highest infectious dose (7.9 × 10^6^ CFU/fish) were euthanized, internal organs were sampled and streaked on TYESA. No bacterial growth occurred from any organ/fish, indicating that the fish had eliminated injected bacteria. These results suggest that this species is non-pathogenic for rainbow trout under the conditions tested. One fish injected with the highest dose (1.9 × 10^6^ CFU) of *F. collinsii* TRV642 died at 12 dpi (Figure 2C) and the bacterium was successfully recovered from internal organs of this fish including spleen, kidney and liver. We also examined at the end of the experiment internal organs of 13 other fish and observed striking differences linked to the fish status. Indeed, starting from 9 dpi, it was noticed in several tanks that dominant fish of bigger size were attacking smaller fish. This behavior can be observed when fish are maintained in small groups, independently of any challenge. The attacked fish then shows heavily damaged caudal fin and caudal tissue erosion (Figure 2D) leading to swimming defects. For ethical reasons, we eliminated the dominant fish by euthanasia starting from 14 dpi and ceased the monitoring of mortality. In some tanks, another fish rapidly replaced the dominant fish and continued to attack weaker fish. Between 14 and 20 dpi, dominant and attacked fish were euthanized and internal organs sampled and streaked on TYESA. Out of 8 attacked fish, *Flavobacterium collinsii* was successfully recovered from internal organs of 7 fish. In sharp contrast, none of the 5 examined dominant fish was positive for *F. collinsii*. Taken together, these results indicate that, in our experimental set-up, *F. collinsii* is able to invade fish with compromised conditions such as stress and/or wounds.

## DISCUSSION

Since its thorough 1996 revision on the basis of 16S DNA sequencing (10), the genus *Flavobacterium* has considerably expanded as a result of the growing number of worldwide sampling campaigns (https://lpsn.dsmz.de/search?word=flavobacterium). Most of the *Flavobacterium* species described so far are *a priori* non-pathogenic, environmental isolates, retrieved from a wide diversity of sources (*e.g*., fresh and salt water, soil, rhizosphere). However, a number of new species have also been described from fish or fish farm environments, which raised questions on whether some could cause disease (42) as *F. columnare* and *F. psychrophilum*: *F. columnare* is the causative agent of columnaris disease which affects a large variety of freshwater fish species generally reared at relatively warm temperatures such as catfish, tilapia and ornamental fish (43); *F. psychrophilum* causes Rainbow Trout Fry Syndrome and Bacterial Cold-Water Disease affecting predominantly salmonid reared in cold freshwater. To a lesser extent, Bacterial Gill Disease elicited by *F. branchiophilum* also affects a number of cultured fish species throughout the world (44). However, this latter species is considered by some authors as an opportunistic pathogen, arising under suboptimal environmental conditions (16). In addition to these species, a bunch of *Flavobacterium* species retrieved from diseased fish have been described. A large number of uncharacterized fish-associated *Flavobacterium spp*. were also identified during an 8-years follow-up study of the Laurentian Great Lakes in Michigan (45). However, although these bacteria were indeed retrieved from diseased fish, it was unclear whether the symptoms could actually be attributed to the bacteria that were recovered from the infected fish. There is a serious lack of studies aiming to demonstrate bacterial virulence using experimental infection models to fulfill Koch’s postulates. It is also unclear which bacteria are true pathogens and which are opportunistic pathogens or even only secondary colonizers, *i.e*., primarily saprophytic or commensal bacteria able to invade the host only when its defenses are compromised (*e.g*., by wounds, stress, or disease). This information is crucial for appropriate management measures to mitigate disease emergence and spreading.

In this study, we report the isolation of *F. collinsii* strain TRV642 from the spleen of a dead rainbow trout raised in an INRAE experimental fish facility. The dead trout did not display any disease symptoms and belonged to a highly disease-susceptible A36 isogenic line that should be considered as an outstandingly fragile (31). The origin of the contamination is questionable. The facility uses chlorinated tap water, filtration on active charcoal and UV treatment. To guarantee bio-secure conditions, the material is frequently subject to disinfection (bleach) and dry-up periods by qualified staff ensuring BSL2 containments. In addition, in order to ensure a Specific Pathogen Free status, only eggs enter the facility after disinfection (Romeiode®) and the presence of *Flavobacterium* species is checked by bacteriological controls in case of any suspicion after the early stages of development.

Whole genome assembly revealed limited differences in genome size between the isolate TRV642 of *F. collinsii* and the type strain CECT 7796^T^. Strikingly, these genome sizes are far above the mean (4.05 Mbp) and median (3.80 Mbp) genome sizes in the genus *Flavobacterium*. With 5.74 Mbp, strain CECT 7796^T^ possesses the 15^th^ largest genome out of 195 available for the genus (Supplemental data set 1). On the other hand, the two unquestionable fish pathogenic species *F. psychrophilum* and *F. columnare* (as well as its newly described sister-species “*F. covae”, “F. davisii”, and “F. oreochromis”*) possess rather compact, about half smaller, genomes. The presumptively opportunistic pathogen *F. branchiophilum* also harbors a relatively small-sized genome of about 3.6 Mbp. The genome of *F. collinsii* is in the same range of size than those of environmental members of the genus, which suggests an extended ecological niche. Indeed, bacteria living in habitats with diversified nutrient supply tend to have larger genomes and target more complex substrates (46), whereas pathogenicity is generally associated with reduced genome size (47).

The vast majority of the *Flavobacterium* species harboring large genomes are grouped in the phylogenetic tree reconstructed on the alignment of the core genome (Figure 1) suggesting a common ancestor with a large genome. Since these species were retrieved from a variety of ecological niches (*e.g*., sea water, fresh-water, soil), it is difficult to formulate a hypothesis for the habitat of this ancestor. Nevertheless, inside or outside this group some subgroup of bacteria isolated from a same type of habitat were identified. In particular, *F. collinsii*, tightly clusters with *F. tructae* and several other species (*i.e*., *F. chilense, F. piscis, F. bizetiae* and *F. araucananum*) considered as putative fish pathogens because isolated from diseased fish. This common source of isolation suggests evolutionary conserved characteristics. Similarly, a number of species from cold environments including *F. glaciei* grouped together elsewhere in the tree.

It is noteworthy that the picture for the habitat of the different species may be blurred by the very limited number of isolates reported for the vast majority of the considered species which lead us to base our classification essentially on the origin of the type strain. Confining a species to a single habitat may be an oversimplifying hypothesis, especially in the case of the heterotrophic bacteria of the *Flavobacterium* genus. For instance, in 2013, Loch et al. (45) described 32 clusters of isolates belonging to the genus *Flavobacterium* recovered from diseased as well as apparently healthy wild, feral, and farmed fish in Michigan, many of which - most likely representing novel species - displayed significant similarities to environmental species such as *F. hercynium, F. pectinovorum* and *F. frigidimaris*. Furthermore, a 2015 study reported that farmed freshwater fish carried *F. suncheonense, F. indicum, F. aquaticum, F. granuli, F. hercynium* and *F. terrae*, all previously described as environmental species (48). These studies highlight the extreme diversity of *Flavobacterium* species retrieved from fish and the difficulty to identify pathogens.

In line with its large genome, *F. collinsii* TRV642 possesses a broad gene repertoire encompassing 29 PULs likely dedicated to the harvesting of nutrients from the extracellular environment. In addition, the diversity of T9SS cargo proteins, including peptidases and CAZymes, together with others enzymes dedicated to carbohydrates gathering and breakdown, suggests metabolic versatility allowing the bacterium to utilize a large spectrum of carbon sources. Indeed, numerous copies of PULs were identified in environmental species of the family *Flavobacteriaceae*, especially those that are able to utilize a large variety of nutrients sources (*e.g*., 20 *susCD*-like pairs in *Gramella forsetii* KT0803, 42 in *F. johnsoniae* UW101 and 71 in *Zobellia galactanivorans* Dsij) whereas very few pairs were identified in the *bona-fide* fish-pathogenic species (*e.g*., only 1 *susCD*-like pair in *F. psychrophilum* JIP02/86, 2 in *F. columnare* ATCC 23463^T^ and 6 in *Tenacibaculum maritimum* NCIMB 2154^T^). With reference to the Brillat-Savarin’s aphorism “Tell me what you eat, and I will tell you what you are”, one should consider that *F. collinsii* and probably most, if not all, the aforementioned species occasionally retrieved from fish and harboring large genomes, are very likely versatile in their diets. As a result, one might suggest that, as reported for others *Bacteroidota*, these bacteria are associated with niches containing diverse nutrient sources, including eukaryotic organisms, with whom they may have evolved a range of relationships potentially from symbiotic, mutualistic or commensal, to pathogenic interactions. The presence of antibiotic biosynthesis gene clusters in the *F. collinsii* genome was also reported in some mutualistic or commensal bacteria where they are used to fight other competitors occupying the same niche (49), a phenomenon participating in competitive exclusion.

Using a rainbow trout experimental infection model based on intramuscular injection, which is generally much more efficient than the immersion model since it bypasses natural protection barriers of mucus and skin, neither mortality nor symptoms were observed using the highest dose (7.9 × 10^6^ CFU/fish) of *F. bernardetii* F-372^T^. Similar conclusion was drown for *F. collinsii* TRV642, except that when aggressive behavior of dominant fish was observed, the bacterium was recovered from various organs (spleen, kidney and liver) of the attacked fish but not from the dominant fish, suggesting that the infection developed better in fish submitted to stresses and/or wounds. It is also noteworthy that *F. collinsii* TRV642 was originally isolated from a highly *F. psychrophilum*-susceptible A36 isogenic rainbow trout line. In contrast to *F. bernardetii* and *F. collinsii*, experimental infection with *F. psychrophilum* of the same Sy rainbow trout line resulted in high mortality: the median lethal doses (LD50) ranged from ~10^5^ CFU for strains with moderate virulence such as OSU THCO2-90 (50) to < 10^3^ CFU for highly virulent strains as documented for FRGDSA 1882/11 in this study. However, we cannot exclude that different rainbow trout genetic backgrounds, rearing conditions, bacterial growth conditions or even other *F. bernardetii* and *F. collinsii* isolates could actually produce disease.

Many members of the family *Flavobacteriaceae*, and more generally of the phylum *Bacteroidota*, are normal constituents of the host microbiota. However, some of their traits could be considered as ‘dual use’ virulence traits because, while serving important functions for survival in the environment, they can also function as virulence factors (51). Bacteria encompassing these ‘dual use’ virulence traits can rapidly proliferate to degrade and metabolize host macromolecules under certain conditions as recently reported for *Bacteroidota* causing opportunistic diseases in marine eukaryotes (52).

Taken together, our results suggest that some recently described *Flavobacterium* species isolated from fish are likely saprophytic or commensal bacteria that may behave as opportunistic pathogens able to proliferate on decaying tissue to exploit the available nutrients, causing disease under specific circumstances. Indeed, commercially reared fish are subjected to intensive farming practices resulting in welfare issues associated with high stocking density and handling processes that can result in trauma providing favorable conditions for bacteria to thrive.

## ACKNOWLEDGEMENTS

This work was financially supported by the Agence Nationale de la Recherche (grant ANR-17-CE20-0020-01 FlavoPatho) and by institutional support from INRAE. The authors are grateful to the staff of the fish facilities (IERP, INRAE, Infectiology of Fishes and Rodent Facility, doi: 10.15454/1.5572427140471238E12; and PEIMA, INRAE, Fish Farming Systems Experimental Facility, doi: 10.15454/1.5572329612068406E12) for supplying fish and for technical assistance and advice. Animal experiments and sampling were performed in accordance with the European Directive 2010/2063/UE, and were approved by the ethics committee of the INRAE Center in Jouy-en-Josas, France (authorization number 2015100215242446). The authors are grateful to the facilities and expertise of the high throughput-sequencing platform of IMAGIF (http://www.i2bc.paris-saclay.fr/), the INRAE MIGALE bioinformatics facility (https://doi.org/10.15454/1.5572390655343293E12), the MicroScope annotation platform and the National Infrastructure ‘France Génomique’ for providing computational resources.

## CONFLICT OF INTEREST STATEMENT

The authors declare no conflict of interest.

## DATA AVAILABILITY

The annotated TRV642 genome sequence has been deposited at ENA under the accession number ERZ13686542.

## REFERENCES

1. McBride MJ. 2014. The Family *Flavobacteriaceae*, p. 643–676. *In* Rosenberg, E, DeLong, EF, Lory, S, Stackebrandt, E, Thompson, F (eds.), The prokaryotes: Other major lineages of bacteria and the archaea. Springer, Berlin, Heidelberg.

2. Thomas F, Hehemann J-H, Rebuffet E, Czjzek M, Michel G. 2011. Environmental and gut *Bacteroidetes*: The food connection. Front Microbiol 2:93.

3. McBride MJ. 2019. *Bacteroidetes* gliding motility and the Type IX Secretion System. Microbiol Spectr 7.

4. LaFrentz BR, Králová S, Burbick CR, Alexander TL, Phillips CW, Griffin MJ, Waldbieser GC, García JC, de Alexandre Sebastião F, Soto E, Loch TP, Liles MR, Snekvik KR. 2022. The fish pathogen *Flavobacterium columnare* represents four distinct species: *Flavobacterium columnare, Flavobacterium covae* sp. nov., *Flavobacterium davisii* sp. nov. and *Flavobacterium oreochromis* sp. nov., and emended description of *Flavobacterium columnare*. Syst Appl Microbiol 45:126293.

5. Kämpfer P, Lodders N, Martin K, Avendaño-Herrera R. 2012. *Flavobacterium chilense* sp. nov. and *Flavobacterium araucananum* sp. nov., isolated from farmed salmonid fish. Int J Syst Evol Microbiol 62:1402–1408.

6. Saticioglu IB, Ay H, Altun S, Sahin N, Duman M. 2021. *Flavobacterium bernardetii* sp. nov., a possible emerging pathogen of farmed rainbow trout (*Oncorhynchus mykiss*) in cold water. Aquaculture 540:736717.

7. Saticioglu IB, Ay H, Altun S, Duman M, Sahin N. 2021. *Flavobacterium turcicum* sp. nov. and *Flavobacterium kayseriense* sp. nov. isolated from farmed rainbow trout in Turkey. Syst Appl Microbiol 44:126186.

8. Zamora L, Vela AI, Sánchez-Porro C, Palacios MA, Domínguez L, Moore ERB, Ventosa A, Fernández-Garayzábal JF. 2013. Characterization of flavobacteria possibly associated with fish and fish farm environment. Description of three novel *Flavobacterium* species: *Flavobacterium collinsii* sp. nov., *Flavobacterium branchiarum* sp. nov., and *Flavobacterium branchiicola* sp. nov. Aquaculture 416–417:346–353.

9. Strohl WR, Tait LR. 1978. *Cytophaga aquatilis* sp. nov., a facultative anaerobe isolated from the gills of freshwater fish. Int J Syst Evol Microbiol 28:293–303.

10. Bernardet J-F, Segers P, Vancanneyt M, Berthe F, Kersters K, Vandamme P. 1996. Cutting a Gordian Knot: Emended classification and description of the genus *Flavobacterium*, emended description of the Family *Flavobacteriaceae*, and proposal of *Flavobacterium hydatis* nom. nov. (Basonym, Cytophaga aquatilis Strohl and Tait 1978). Int J Syst Evol Microbiol, 46:128–148.

11. de Alexandre Sebastião F, LaFrentz BR, Shelley JP, Stevens B, Marancik D, Dunker F, Reavill D, Soto E. 2019. *Flavobacterium inkyongense* isolated from ornamental cichlids. J Fish Dis 42:1309–1313.

12. Flemming L, Rawlings D, Chenia H. 2007. Phenotypic and molecular characterisation of fish-borne *Flavobacterium johnsoniae*-like isolates from aquaculture systems in South Africa. Res Microbiol 158:18–30.

13. Zamora L, Fernández-Garayzábal JF, Svensson-Stadler LA, Palacios MA, Domínguez L, Moore ERB, Vela AI. 2012. *Flavobacterium oncorhynchi* sp. nov., a new species isolated from rainbow trout (*Oncorhynchus mykiss*). Syst Appl Microbiol 35:86–91.

14. Zamora L, Vela AI, Sánchez-Porro C, Palacios MA, Moore ERB, Domínguez L, Ventosa A, Fernández-Garayzábal JF. 2014. *Flavobacterium tructae* sp. nov. and *Flavobacterium piscis* sp. nov., isolated from farmed rainbow trout (*Oncorhynchus mykiss*). Int J Syst Evol Microbiol 64:392–399.

15. Zamora L, Fernández-Garayzábal JF, Sánchez-Porro C, Palacios MA, Moore ERB, Domínguez L, Ventosa A, Vela AI. 2013. *Flavobacterium plurextorum* sp. nov. isolated from farmed rainbow trout (*Oncorhynchus mykiss*). PLoS One 8:e67741.

16. Good C, Davidson J, Wiens GD, Welch TJ, Summerfelt S. 2015. *Flavobacterium branchiophilum* and *F. succinicans* associated with bacterial gill disease in rainbow trout *Oncorhynchus mykiss* (Walbaum) in water recirculation aquaculture systems. J Fish Dis 38:409–413.

17. Loch TP, Faisal M. 2014. *Flavobacterium spartansii* sp. nov., a pathogen of fishes, and emended descriptions of *Flavobacterium aquidurense* and *Flavobacterium araucananum*. Int J Syst Evol Microbiol 64:406–412.

18. Loch TP, Faisal M. 2016. *Flavobacterium spartansii* induces pathological changes and mortality in experimentally challenged Chinook salmon *Oncorhynchus tshawytscha* (Walbaum). J Fish Dis 39:483–488.

19. D’Ambrosio J, Phocas F, Haffray P, Bestin A, Brard-Fudulea S, Poncet C, Quillet E, Dechamp N, Fraslin C, Charles M, Dupont-Nivet M. 2019. Genome-wide estimates of genetic diversity, inbreeding and effective size of experimental and commercial rainbow trout lines undergoing selective breeding. Genetics Selection Evolution 51:26.

20. Thompson WR. 1947. Use of moving averages and interpolation to estimate median-effective dose. Bacteriological Reviews 11:115–145.

21. Davis JJ, Wattam AR, Aziz RK, Brettin T, Butler R, Butler RM, Chlenski P, Conrad N, Dickerman A, Dietrich EM, Gabbard JL, Gerdes S, Guard A, Kenyon RW, Machi D, Mao C, Murphy-Olson D, Nguyen M, Nordberg EK, Olsen GJ, Olson RD, Overbeek JC, Overbeek R, Parrello B, Pusch GD, Shukla M, Thomas C, VanOeffelen M, Vonstein V, Warren AS, Xia F, Xie D, Yoo H, Stevens R. 2020. The PATRIC Bioinformatics Resource Center: expanding data and analysis capabilities. Nucleic Acids Res 48:D606–D612.

22. Vallenet D, Calteau A, Dubois M, Amours P, Bazin A, Beuvin M, Burlot L, Bussell X, Fouteau S, Gautreau G, Lajus A, Langlois J, Planel R, Roche D, Rollin J, Rouy Z, Sabatet V, Médigue C. 2020. MicroScope: an integrated platform for the annotation and exploration of microbial gene functions through genomic, pangenomic and metabolic comparative analysis. Nucleic Acids Res 48:D579–D589.

23. Haft DH, Selengut JD, White O. 2003. The TIGRFAMs database of protein families. Nucleic Acids Res 31:371–373.

24. Huang L, Zhang H, Wu P, Entwistle S, Li X, Yohe T, Yi H, Yang Z, Yin Y. 2018. dbCAN-seq: a database of carbohydrate-active enzyme (CAZyme) sequence and annotation. Nucleic Acids Res 46:D516–D521.

25. Rawlings ND, Barrett AJ, Thomas PD, Huang X, Bateman A, Finn RD. 2018. The MEROPS database of proteolytic enzymes, their substrates and inhibitors in 2017 and a comparison with peptidases in the PANTHER database. Nucleic Acids Res 46:D624–D632.

26. Yoon S-H, Ha S, Lim J, Kwon S, Chun J. 2017. A large-scale evaluation of algorithms to calculate average nucleotide identity. Antonie van Leeuwenhoek 110:1281–1286.

27. O’Leary NA, Wright MW, Brister JR, Ciufo S, Haddad D, McVeigh R, Rajput B, Robbertse B, Smith-White B, Ako-Adjei D, Astashyn A, Badretdin A, Bao Y, Blinkova O, Brover V, Chetvernin V, Choi J, Cox E, Ermolaeva O, Farrell CM, Goldfarb T, Gupta T, Haft D, Hatcher E, Hlavina W, Joardar VS, Kodali VK, Li W, Maglott D, Masterson P, McGarvey KM, Murphy MR, O’Neill K, Pujar S, Rangwala SH, Rausch D, Riddick LD, Schoch C, Shkeda A, Storz SS, Sun H, Thibaud-Nissen F, Tolstoy I, Tully RE, Vatsan AR, Wallin C, Webb D, Wu W, Landrum MJ, Kimchi A, Tatusova T, DiCuccio M, Kitts P, Murphy TD, Pruitt KD. 2016. Reference sequence (RefSeq) database at NCBI: current status, taxonomic expansion, and functional annotation. Nucleic Acids Res 44:D733–745.

28. Camacho C, Coulouris G, Avagyan V, Ma N, Papadopoulos J, Bealer K, Madden TL. 2009. BLAST+: architecture and applications. BMC Bioinformatics 10:421.

29. Edgar RC. 2004. MUSCLE: multiple sequence alignment with high accuracy and high throughput. Nucleic Acids Res 32:1792–1797.

30. Price MN, Dehal PS, Arkin AP. 2009. FastTree: computing large minimum evolution trees with profiles instead of a distance matrix. Mol Biol Evol 26:1641–1650.

31. Fraslin C, Quillet E, Rochat T, Dechamp N, Bernardet J-F, Collet B, Lallias D, Boudinot P. 2020. Combining multiple approaches and models to dissect the genetic architecture of resistance to infections in fish. Front Genet 11:677.

32. Richter M, Rosselló-Móra R. 2009. Shifting the genomic gold standard for the prokaryotic species definition. PNAS 106:19126–19131.

33. Mühle E, Abry C, Leclerc P, Goly G-M, Criscuolo A, Busse H-J, Kämpfer P, Bernardet J-F, Clermont D, Chesneau O 2021. 2020. *Flavobacterium bizetiae* sp. nov., isolated from diseased freshwater fish in Canada at the end of the 1970s. Int J Syst Evol Microbiol 71:004576.

34. McBride MJ, Xie G, Martens EC, Lapidus A, Henrissat B, Rhodes RG, Goltsman E, Wang W, Xu J, Hunnicutt DW, Staroscik AM, Hoover TR, Cheng Y-Q, Stein JL. 2009. Novel Features of the polysaccharide-digesting gliding bacterium *Flavobacterium johnsoniae* as revealed by genome sequence analysis. Appl Environ Microbiol 75:6864–6875.

35. Duchaud E, Rochat T, Habib C, Barbier P, Loux V, Guérin C, Dalsgaard I, Madsen L, Nilsen H, Sundell K, Wiklund T, Strepparava N, Wahli T, Caburlotto G, Manfrin A, Wiens GD, Fujiwara-Nagata E, Avendaño-Herrera R, Bernardet J-F, Nicolas P. 2018. Genomic diversity and evolution of the fish pathogen *Flavobacterium psychrophilum*. Front Microbiol 9:138.

36. Li YG, Hu B, Christie PJ. 2019. Biological and structural diversity of Type IV Secretion Systems. Microbiol Spectr 7.

37. Pollet RM, Martin LM, Koropatkin NM. 2021. TonB-dependent transporters in the *Bacteroidetes*: Unique domain structures and potential functions. Mol Microbiol 115:490–501.

38. Terrapon N, Lombard V, Gilbert HJ, Henrissat B. 2015. Automatic prediction of polysaccharide utilization loci in *Bacteroidetes* species. Bioinformatics 31:647–655.

39. Glenwright AJ, Pothula KR, Bhamidimarri SP, Chorev DS, Baslé A, Firbank SJ, Zheng H, Robinson CV, Winterhalter M, Kleinekathöfer U, Bolam DN, van den Berg B. 2017. Structural basis for nutrient acquisition by dominant members of the human gut microbiota. 7637. Nature 541:407–411.

40. Madej M, White JBR, Nowakowska Z, Rawson S, Scavenius C, Enghild JJ, Bereta GP, Pothula K, Kleinekathoefer U, Baslé A, Ranson NA, Potempa J, van den Berg B. 2020. Structural and functional insights into oligopeptide acquisition by the RagAB transporter from *Porphyromonas gingivalis*. Nat Microbiol 5:1016–1025.

41. Su H, Blain F, Musil RA, Zimmermann JJ, Gu K, Bennett DC. 1996. Isolation and expression in *Escherichia coli* of hepB and hepC, genes coding for the glycosaminoglycan-degrading enzymes heparinase II and heparinase III, respectively, from *Flavobacterium heparinum*. Appl Environ Microbiol 62:2723–2734.

42. Loch TP, Faisal M. 2015. Emerging flavobacterial infections in fish: A review. J Adv Res 6:283.

43. Declercq AM, Haesebrouck F, Van den Broeck W, Bossier P, Decostere A. 2013. Columnaris disease in fish: a review with emphasis on bacterium-host interactions. Vet Res 44:27.

44. Ostland VE, MacPHEE DD, Lumsden JS, Ferguson HW. 1995. Virulence of *Flavobacterium branchiophilum* in experimentally infected salmonids. Journal of Fish Diseases 18:249–262.

45. Loch TP, Fujimoto M, Woodiga SA, Walker ED, Marsh TL, Faisal M. 2013. Diversity of fish-associated flavobacteria of Michigan. Journal of Aquatic Animal Health 25:149–164.

46. Grondin JM, Tamura K, Déjean G, Abbott DW, Brumer H. 2017. Polysaccharide Utilization Loci: Fueling Microbial Communities. J Bacteriol 199:e00860–16.

47. Murray GGR, Charlesworth J, Miller EL, Casey MJ, Lloyd CT, Gottschalk M, Tucker AW (Dan), Welch JJ, Weinert LA. 2021. Genome reduction is associated with bacterial pathogenicity across different scales of temporal and ecological divergence. Molecular Biology and Evolution 38:1570–1579.

48. Verma DK, Rathore G. 2015. New host record of five *Flavobacterium* species associated with tropical fresh water farmed fishes from North India. Braz J Microbiol 46:969–976.

49. van der Meij A, Worsley SF, Hutchings MI, van Wezel GP. 2017. Chemical ecology of antibiotic production by actinomycetes. FEMS Microbiology Reviews 41:392–416.

50. Zhu Y, Lechardeur D, Bernardet J-F, Kerouault B, Guérin C, Rigaudeau D, Nicolas P, Duchaud E, Rochat T. 2022. Two functionally distinct heme/iron transport systems are virulence determinants of the fish pathogen *Flavobacterium psychrophilum*. Virulence 13:1221–1241.

51. Casadevall A, Steenbergen JN, Nosanchuk JD. 2003. ‘Ready made’ virulence and ‘dual use’ virulence factors in pathogenic environmental fungi — the *Cryptococcus neoformans* paradigm. Curr Opin Microbiol 6:332–337.

52. Hudson J, Egan S. 2022. Opportunistic diseases in marine eukaryotes: Could *Bacteroidota* be the next threat to ocean life? Environ Microbiol https://doi.org/10.1111/1462-2920.16094.

